# Artifacts in EEG-based BCI therapies: friend or foe?

**DOI:** 10.1101/2021.10.27.466131

**Authors:** Eric James McDermott, Philipp Raggam, Sven Kirsch, Paolo Belardinelli, Ulf Ziemann, Christoph Zrenner

## Abstract

EEG-based brain-computer interfaces (BCI) have promising therapeutic potential beyond traditional neurofeedback training, such as enabling personalized and optimized virtual reality (VR) neurorehabilitation paradigms where the timing and parameters of the visual experience is synchronized with specific brain-states. While BCI algorithms are often designed to focus on whichever portion of a signal is most informative, in these brain-state-synchronized applications, it is of critical importance that the resulting decoder is sensitive to physiological brain activity representative of various mental states, and not to artifacts, such as those arising from naturalistic movements. In this study, we compare the relative classification accuracy with which different motor tasks can be decoded from both extracted brain activity and artifacts contained in the EEG signal. EEG data was collected from 17 chronic stroke patients while performing six different head, hand, and arm movements in a realistic VR-based neurorehabilitation paradigm. Results show that the artifactual component of the EEG signal is significantly more informative than brain activity with respect to classification accuracy. This finding is consistent across different feature extraction methods and classification pipelines. While informative brain signals can be recovered with suitable cleaning procedures, we recommend that features should not be designed solely to maximize classification accuracy, as this could select for remaining artifactual components. We also propose the use of machine learning approaches that are interpretable to verify that classification is driven by physiological brain-states. In summary, whereas informative artifacts are a helpful friend in BCI-based communication applications, they can be a problematic foe in the estimation of physiological brain states.

## 1. Introduction

### 1.1 Motivation

Brain-computer interfaces (BCI) are becoming increasingly applied in rehabilitative settings. At the root of every BCI is the transformation of recorded activity into quantifiable outputs. Yet, brain activity, recorded as data measured from electroencephalogram (EEG), is in general mixed with artifacts such as those arising from muscle activity during the same time period [1]. Since the voltage potentials of muscle activity measured with surface electrodes are several orders of magnitude higher than those generated by brain activity, this can cause BCI algorithms to learn to generate optimal output based on artifacts [2]. While this may be acceptable (and even increase accuracy rates) in the use of BCI as an actuator (for example, in letter selection or wheelchair operation), the classification result is then not a reflection of high-level neuronal brain state but rather of concurrently generated artifacts.

The mixing of brain signals and artifacts can be problematic in BCI applications designed to detect a specific brain state, such as in personalized neurorehabilitation [3]. Therefore, in these cases, it is essential to first distinguish between brain and artifact, especially when decoding brain activity during movement execution. The problem in distinguishing movement related artifacts from movement related neuronal activity is precisely that the artifacts are not random noise: They contaminate the signal of interest (i.e. the higher-level brain state) in a predictable way, and may thereby be even more informative from the point of view of an automated classifier. We therefore consider it relevant to compare classification accuracy from the artifact components of the EEG signal in relation to brain signal components. In this study, we (1) characterize movement artifacts from different movement primitives, (2) separate data into brain signal and muscle- and eye-based artifact signal, (3) use machine learning classifiers to predict movement from both brain and artifact components, and (4) interpret the results to understand what features the classifier identifies as informative.

### 1.2 EEG Artifacts and Processing Pipelines

The most obvious artifacts arise from muscle activity or eye movements, however, cardiac and sweat-related artifacts [4], as well as 50/60 Hz power line noise [5] also play a relevant role. Even though the application of EEG in rehabilitation paradigms is expanding, there is no generally consensus on the procedure for dealing with artifacts, especially those arising from naturalistic movements during the EEG recording. Typically, processing pipelines include a step to remove bad channels and then perform either principal component analysis (PCA) or independent component analysis (ICA) with the components selected manually by visual inspection [2, 6-8]. However, important methodological details are often omitted in literature [1, 2, 8, 9].

A typical subsequent step in standard state-of-the-art EEG signal classification pipelines is to use spatial filters to increase class separability [10]. One of the most robust and effective methods is the common spatial pattern (CSP) algorithm, which finds spatial pattern projections that maximize the variance between classes [11, 12]. However, since this method blindly maximizes separation, it could be susceptible to maximizing the importance of co-occurring artifacts. Apart from CSP, a large variety of different EEG preprocessing pipelines have been put forward using different parameters [13, 14] and it has become a separate area of study to compare the accuracy of automated detection algorithms.

In general, artifacts are considered in the literature to be detrimental to classification accuracy [1], even though their advantage in special cases is acknowledged [15, 16], such as the use of eye blinks or facial muscles to control computers. Whether or not EEG artifacts are problematic in BCI-based neurorehabilitation during naturalistic movements is an open question that we address in this study.

### 1.3 Study Design

BCIs in neurorehabilitation are typically related to recovering a specific lost or impaired function (see a recent review for extensive background information [17]. We therefore designed our paradigm to test the influence of physiologically relevant movements that are frequently impaired by stroke on the EEG signal. Our participants were guided through a virtual reality (VR) paradigm, in which they observed the task environment and their arms from a first-person point-of-view perspective. We chose a VR-based paradigm because the combination of EEG and VR allows for a future “closed-loop” application where the EEG-signal influences the VR-paradigm in real-time to optimize treatment outcome. Additionally, VR paradigms are also reported to be more engaging, motivating, and fun than their traditional therapeutic counterparts [18, 19], and we have successfully created a similar closed-loop system using EMG [20].

In order to investigate the relative contribution of brain signal vs. artifact signal to decoding accuracy, we separate the EEG signal into an artifact part and a brain activity part using ICA with an automated algorithm. We then test how well the specific naturalistic movement that occurred during the respective trial is predicted from each set of data. This is done with two different feature extraction methods (one quantifying average spectral power per channel, and the other considering the time-course of the activity) and two multiclass linear machine learning classifiers. Simultaneous to the EEG recording, we also recorded electromyography (EMG) from the upper limb and neck muscles as a “benchmark” for the classification accuracy that can be achieved from the muscle activity during the movement.

## 2. Materials and Methods

### 2.1 Participants

The study protocol was approved by the local Ethics Review Committee of the Medical Faculty of Eberhard Karls University Tübingen (Protocol BNP-2019-11). The study was conducted in accordance with the latest version of the Declaration of Helsinki. After giving written informed consent, 17 patients who had previously been diagnosed with stroke were included in the study fulfilling the following pre-established inclusion criteria: (i) age of 18–80 years, (ii) participant had an ischemic stroke more than 12 months ago, (iii) participant has a motor impairment of the arm and/or hand as a result of the stroke (iv) participant is otherwise in good physical and mental health.

### 2.2 Experimental Set-up

Scalp EEG was recorded using a 64-channel EEG cap (Easycap GmbH, Munich) in a 10-5 system layout [21] with an additional concentration of electrodes over the motor cortex (see Appendix, Figure A1). Muscle activity was recorded using 7 bipolar surface EMG electrodes (Kendall): Five electrodes were placed on the arm used in the task on the brachioradialis, extensor digitorum, flexor digitorum profundus, biceps, and deltoid, and two more electrodes were placed on the left and right sternocleidomastoid. EEG and EMG signals were acquired simultaneously using a biosignal amplifier (NeurOne Tesla, Bittium, Finland) at a sample rate of 1 kHz in DC. Participants were seated in a comfortable chair, while visual stimulation was provided using the HTC Vive Virtual Reality Headset^1^. Hand positioning and movement was measured using the Valve Index Knuckle Controllers^2^. Timestamps and event triggers were sent into the NeurOne data stream through a user data protocol (UDP) at the start and end of the reference phase, wait period, and task. Synchronization between the task and EMG/EEG activity was achieved through timestamp alignment.

### 2.3 Virtual Reality Presentation

Tasks were implemented using Unreal Engine 4^3^, and were presented to the participant through the VR headset. Prior to beginning the experiment, a proper fit of the headset was achieved for each patient, and verbal confirmation of a “clear image” was obtained. Hand position calibration was performed before the experimental session began by establishing a comfortable position on the chair of the arm, thereby allowing each task to be presented within the patient’s reach.

### 2.4 Task Protocol

Two 3-minute eyes-open resting-state EEG measurements were recorded in sequence, with and without wearing the VR headset, in order to visually inspect the data quality. Patients were then given written instructions accompanied by verbal explanations on how to perform each task. Then, a practice round was carried out, where patients performed each task under verbal guidance until executed correctly. Six different tasks in total were performed, requiring the execution of a particular movement sequence. Each task began with a 2-second fixation phase, where the respective task was indicated by the virtual environment, and during which patients moved toward the starting positions. A 4-second preparation phase followed, which required the patient to maintain a steady head and hand position at the starting position. This 4-second countdown was programmed to automatically restart if any movement was detected during the fixation phase. Once the fixation phase was completed, the patient was free to perform the task without a time limit. A 2-second rest phase then followed, after which the next task was initiated, beginning again with the fixation phase. The lamp task was not included in subsequent analysis due to the trial duration of a button press being too short. The remaining five tasks were used for EEG-classification. Representations of each task can be seen below in Figure 1.

**Figure 1.**
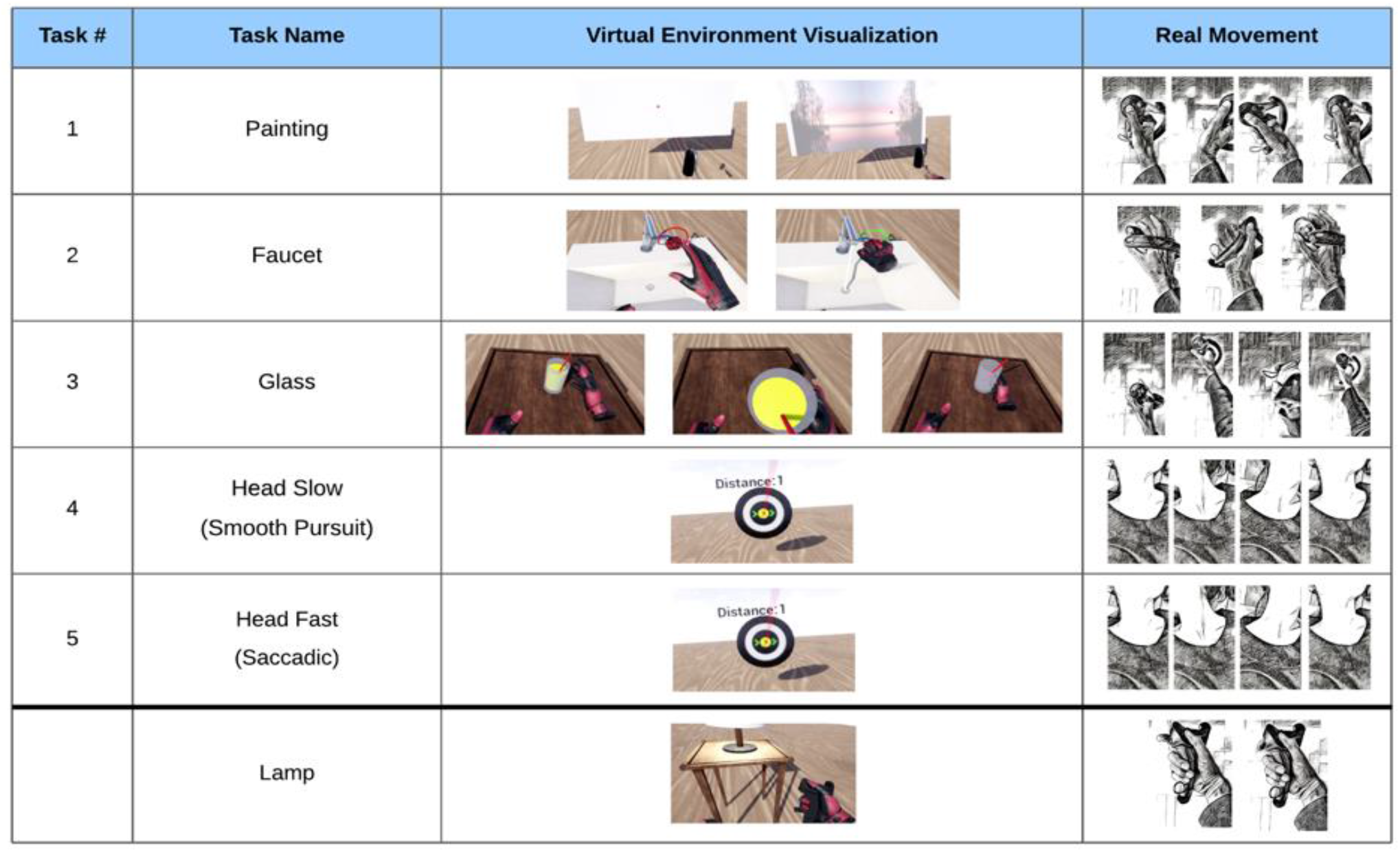
The five tasks with VR visualization and movement depiction. (**Painting**) Wrist extension, followed by flexion, followed by extension. (**Faucet**) Forearm supination of >20°, followed by a pronation. (**Glass**) A complex movement consisting of an elbow extension, a grasp, elbow flexion, elbow extension, and a release of the grasp. (**Head Slow**) Smooth pursuit of the head while tracking a target moving to the right, then left, then right. (**Head Fast**) Saccadic movement of the head toward a target appearing to the right, then left, then right. (**Lamp**) Button press with the index finger or thumb (only included in EMG analysis).

These phases taken together represent the process for a single trial (see Figure 2). In a single run, 10 trials of each task movement were required, followed by a break. Study participants performed 3 runs, fewer if they found it too strenuous. Tasks were presented in random order, with the constraints that the same task would not appear more than twice in a row.

**Figure 2.**
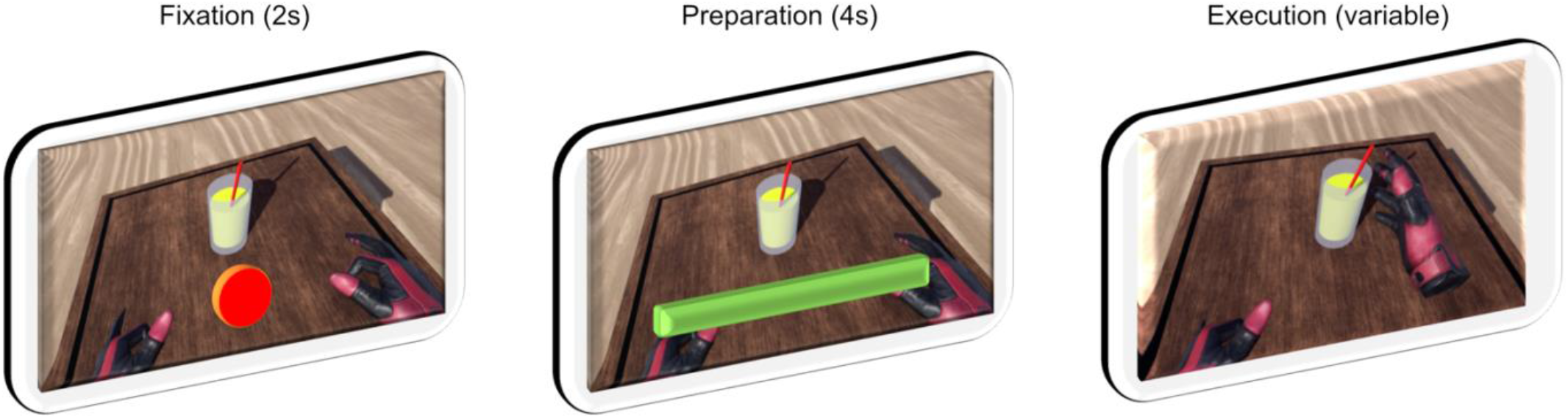
Visualization of one complete trial, along with timings. Rest phase between trials not shown.

### 2.5 Data Preprocessing

Data processing was performed using custom scripts in MATLAB^4^ (R2020a), using EEGLab Toolbox 2020_0 [22], and FastICA toolbox v. 2.5 [23]. EEG and EMG data were down-sampled to 500Hz, high-pass filtered with a cut-off frequency of 0.5Hz and notch-filtered to attenuate 50 Hz electrical line noise. The data was then epoched according to the triggers sent during the tasks. In particular, for each single trial, a *preparation epoch* was extracted between onset of the fixation timer and the onset of the task, and an *execution epoch* was extracted between the task onset and completion. Checks were also performed to ensure that no false triggers were present and/or used. Finally, data was split into EMG and EEG, as seen by the full pipeline visualized in Figure 3.

**Figure 3.**
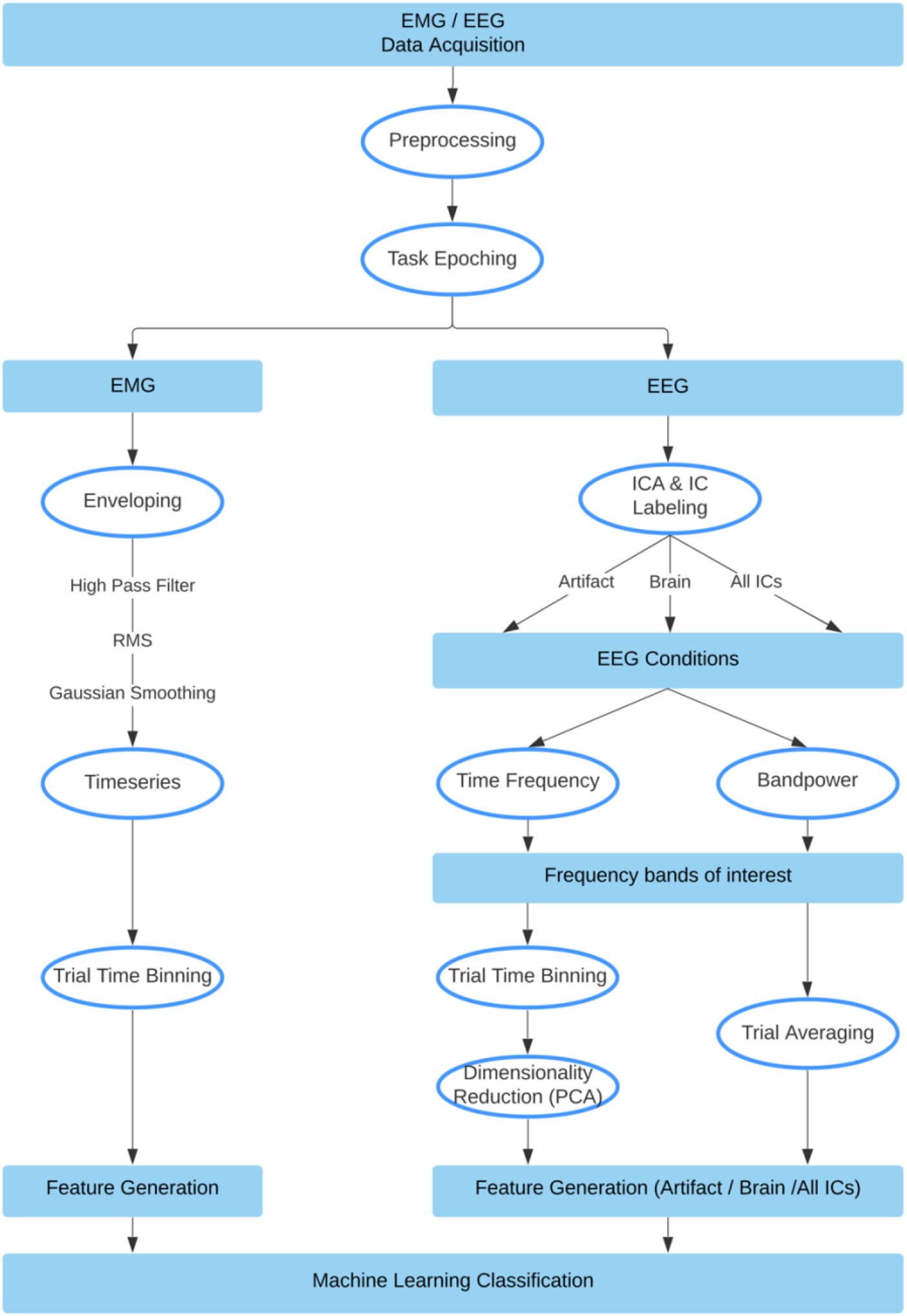
Method Pipeline, split into EMG and EEG pathways.

#### 2.5.1 EMG Data

The resulting epochs were of differing lengths, due to variance in the duration of movement execution. EMG data consisted of 7 bipolar channel recordings, 5 ‘arm’ electrodes and 2 ‘neck’ electrodes. To prepare features from the EMG data for the machine learning classifier task, the data was first high-passed filtered with a cut-off frequency of 10Hz. Then, the envelope of the data was computed using a root mean square (RMS) sliding window of 250ms. This envelope was then gaussian smoothed with a 100ms sliding window. Next, the duration of each envelope was normalized and resampled to contain 1000 time points. This data was then binned by averaging over every 100 time points for each trial, creating a vector containing 10 elements. This procedure was performed for each channel, resulting in 70 feature vectors per participant, per trial; which was used for subsequent classification. The process is visualized in Figure 4.

**Figure 4.**
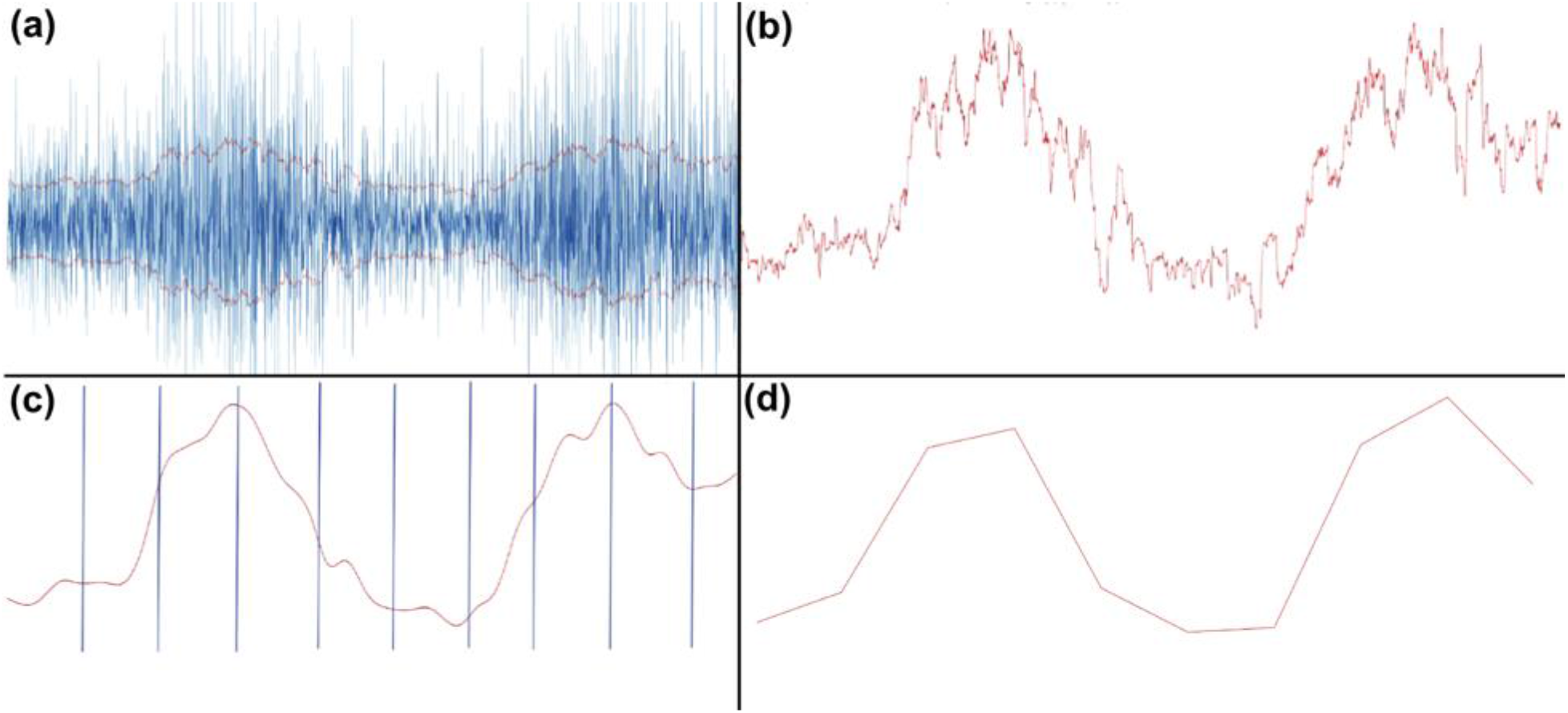
EMG processing steps. (**a**) RMS of high-passed EMG data. (**b**) Positive portion of the envelope. (**c**) Gaussian smoothed and broken into equal 100-point time bins. (**d**) Averaged time bins, resulting in 10 values, plotted here as a line.

#### 2.5.2 EEG Data

Given the nature of a study examining artifacts, we purposefully chose not to remove any of the trials or electrodes in our preprocessing pipeline. EEG data were down-sampled to 500Hz, and then high-pass filtered with a cut-off frequency of 0.5Hz. The data was then notch filtered and then re-referenced to a common average reference. The EEG data was further processed by a baselining in the form of subtracting the mean from each trial epoch. The next step involved performing ICA over each participant’s complete set of task execution data (aggregated across all tasks). We then used an automated process (EEGLAB’s *IClabel* function) to classify the resulting independent components. Components were ranked according to type (Brain, Muscle, Eye, Artifact, Cardiac, Other), and in order of their variance. Using *IClabel*, we selected the top-10 ranked ‘brain’ independent components, as well as the top-10 ranked ‘artifact’ (combined ‘Muscle’ and ‘Eye’) independent components to extract the respective signal portions. This resulted in our 3 EEG conditions: brain-only, artifact-only, and all.

At this point, our EEG processing pipeline split into two approaches: *The Bandpower Approach* and *The Time-Frequency Approach*.

#### 2.5.3 The Time-Frequency Approach

For each condition separately, time frequency analysis (TFA) was performed using FFT in the range of 3-40Hz across each participant, task, and individual trial. We selected 5 different frequency bands: theta (3-7Hz), alpha (7-13Hz), low beta (13-16Hz), beta (16-26Hz), and gamma (26-40Hz), and then averaged across each of these ranges for all trials within a task for each participant. In the brain and artifact conditions, this trial-averaged TF-data was then binned to create 10 time bins for each of the 5 frequency bands for each of the 10 components over each task and each participant, creating 500 (10 × 5 × 10) features for each participant. Whereas in the ‘all’ condition, all 64 components remained, creating 3200 (10 × 5 × 64) features for each participant. To reduce the risk of overfitting, the dimensionality of the data for each condition was reduced to 70 features (to include 14 features for each of the 5 frequency bands) using PCA separately for each task and participant. This data was then passed to the classifier to obtain prediction accuracy. We were also able to examine the classification results of each frequency band independently.

#### 2.5.4 The Bandpower Approach

For each condition separately, the EEG signals were bandpass-filtered in each of six frequency bands: theta (3-7Hz), alpha (7-13Hz), low beta (13-16Hz), beta (16-26Hz), gamma (26-40Hz), and “all” (3-40Hz). The channel-specific bandpower for every single trial was calculated for the band-passed data using the equation:

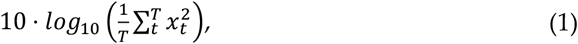

where *x*_*t*_ represents the task signal of a single trial at time-point *t* with the length *T*, over each participant and task. These single values were then the features, resulting in 64 features in total (one for each channel) for each trial and each task. Likewise, we were also able to obtain classification accuracy for each frequency band. To obtain topographical plots, we averaged the trials for each channel.

#### 2.5.5 Common Spatial Patterns (CSP)

The CSP algorithm uses spatial pattern projections to maximize the discriminability between classes. CSP is in general designed for a two-class problem. Here, the MNE^5^ decoding package CSP was used and adapted for the current study’s multiclass-problem [24]. The covariance matrix of the epoched EEG signals from the classes were calculated and sorted by their eigenvalues, in order to find the projections with the highest variances between the classes. For the covariance matrix calculation, the Ledoit-Wolf shrinkage estimator was applied. The number of components (i.e. spatial projections) chosen was four, resulting in a feature reduction from 64 to four. These components were then plotted as topographies and visually compared to the “brain”-only derived independent components. The goal of implementing the CSP algorithm was to inspect the spatial projections (i.e. components) of cleaned EEG signals for artifacts, hence it was only applied to the “brain”-only derived independent components.

### 2.6 Classifier Properties

#### 2.6.1 The Time-Frequency Approach

We input the feature vectors into MATLABs fitcecoc^6^, which is a “*multiclass error-correcting output codes (ECOC)*’’ model that inherently takes a multiclass problem and reduces it to a set of binary learners through the use of support vector machines (SVM). ECOC models have shown to improve accuracy with respect to other multiclass models [25]. In our classifier, we chose to take an all-versus-one approach instead of one-vs-one approach. In general, 70 features were used for the TFA approach and EMG, and 14 features when examining individual frequency bands from the TFA approach. No hyperparameter optimization was performed, as this was not the aim of the current study. For an example of a VR-EEG optimized signal analysis pipeline, please see [3]. The classifier pipeline used an 80-20% train-test split, where the model variables were created in the training phase, and then never-before-seen testing data was used for the testing accuracy. We bootstrapped our model 100 times, using a new 80-20% split of the data each time.

#### 2.6.2 The Bandpower Approach

For this approach, the classification of the tasks was performed in Python^7^, using the *scikit-learn* package^8^. The resulting 64 features were fed into a linear classifier (multiclass linear discriminant analysis, LDA) that used a singular value decomposition solver, which is recommended for data with a large number of features. For the classification, a 10-times 10-fold cross-validation (CV) approach was used. In CV, the dataset was split into 10 pieces, using each piece once as test data and the other 9 pieces as training data (90-10% split), and repeating that procedure 10 times, which resulted in 100 classification accuracy values per participant and frequency band. Participants with less than 10 trials in a condition were excluded for both approaches (2 participants).

## 3 Results

### 3.1 Summary

EEG data recorded during the execution of 5 different movements was demixed into an artifact portion as well as a brain signal component portion using ICA. We then compared how informative each dataset is with regard to predicting (post-hoc) the movement that had been executed during the respective trial. Two different methods were used for feature extraction (time-frequency and bandpower analysis). The data consisting of artifact components was consistently more predictive than the data consisting of brain-signal components. EMG data was collected to serve as a benchmark for the classification accuracy that can be achieved from direct measures of muscle activity.

### 3.2 EMG Analysis

As muscle activity greatly influences EEG during movement, EMG data was recorded as a benchmark for how informative pure muscle activity is for movement classification. Preprocessing of EMG data resulted in distinct patterns of activation for each of the movements that can be visualized by plotting the average rate of change of muscle activity during task execution. Two examples of how the movement sequence maps to a corresponding “*EMG fingerprint*” are shown in Figure 5 (see Appendix, Figure 2A for complete participant data).

**Figure 5.**
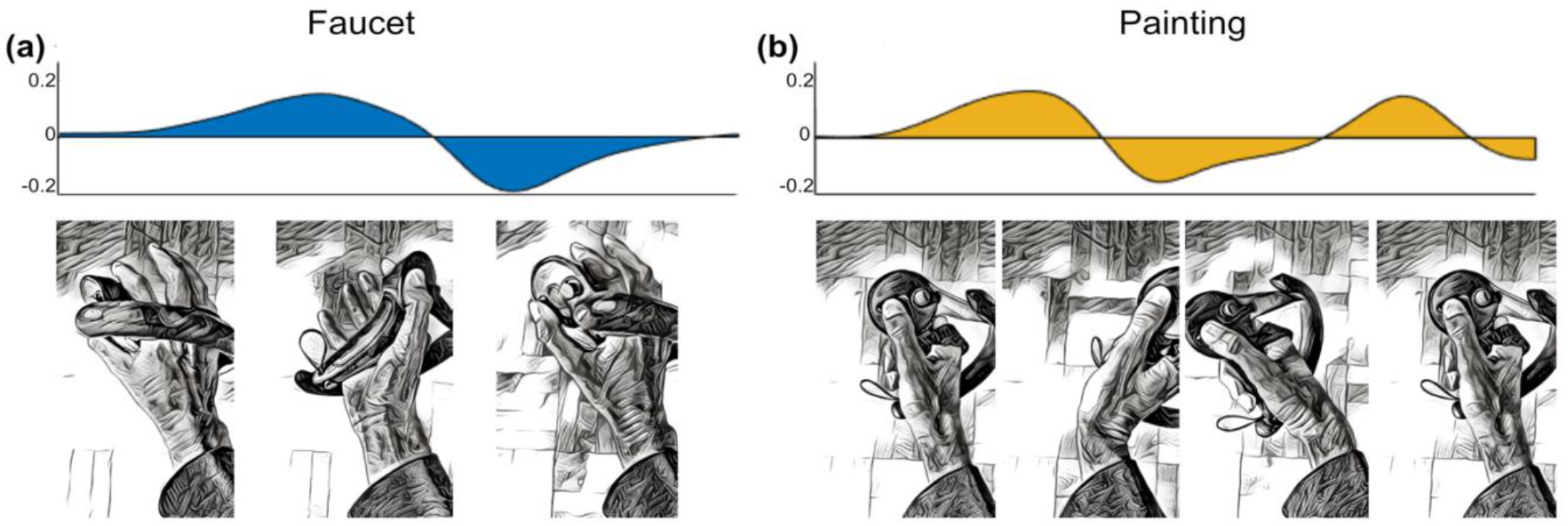
Visualization of EMG activity during task execution (duration normalized) showing the average rate of change of EMG activity across all study participants, arbitrary units. (**a**) Biceps muscle activity during faucet rotation task. (**b**) Extensor digitorum muscle activity during back-and-forth painting task. Hand positions at different phases of the task illustrated below.

### 3.3. Artifact Characterization

Relevant sources of artifacts with respect to BCI signal processing are eye blinks, eye movements and tonic or phasic muscle activity. Representative examples of artifact ICA component topographies are shown in Figure 6, as visualized using EEGLAB’s *IClabel* extension.

**Figure 6.**
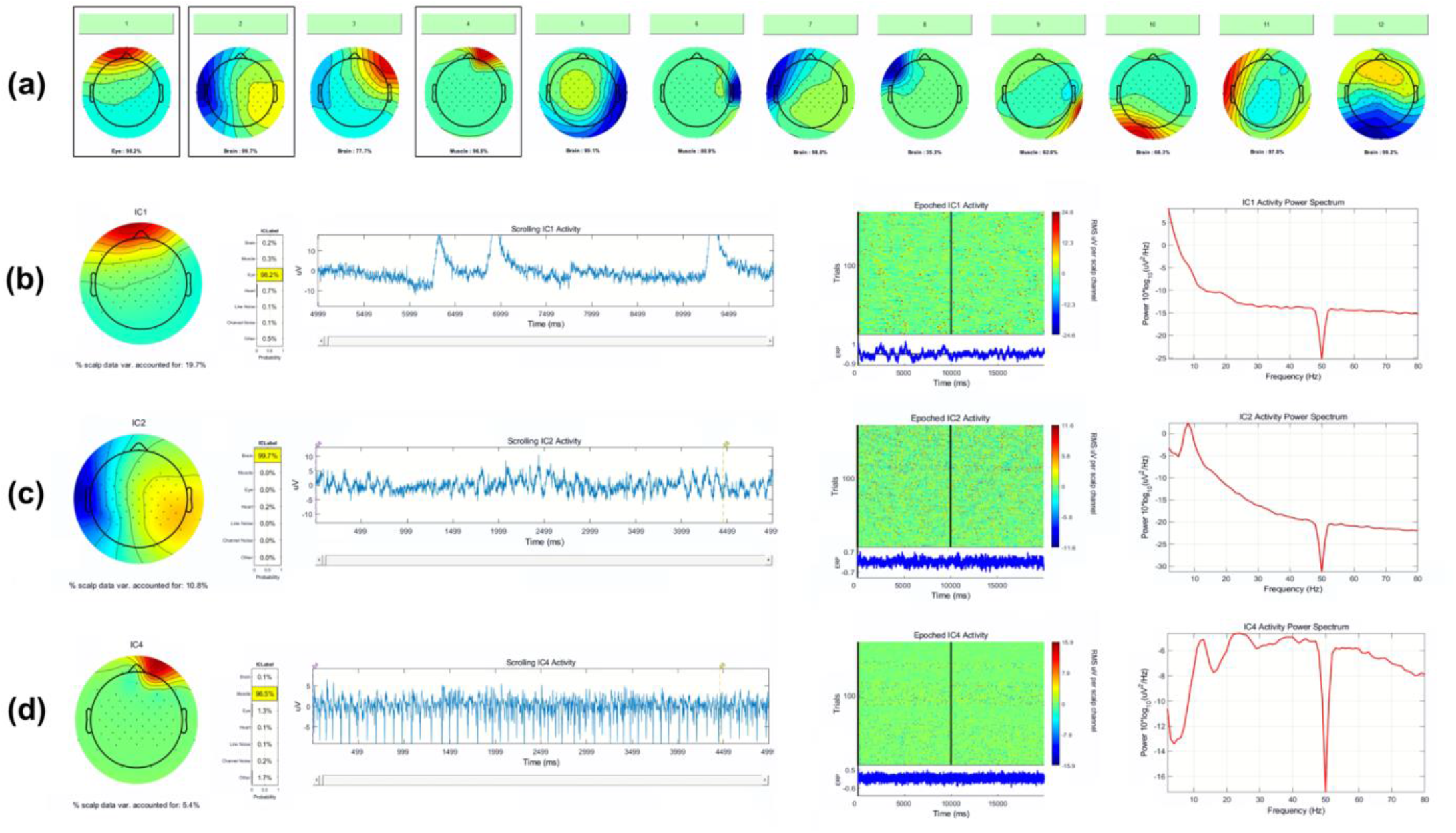
(**a**) Example output from EEGLABs *IClabel* function, indicating the probability of each IC’s class. Here, ICA components 1, 4, 6, and 9 are classified as eye and muscle activity, whereas 2, 3, 5, 7, 8, 10, 11, and 12 are classified as brain activity. Profiles of outlined ICs 1, 2, and 4 are shown in detail. (**b**) Detailed view of IC 1, eye. (**c**) Detailed view of IC 2, brain. (**d**) Detailed view of IC 4, muscle.

### 3.4 Classification Accuracy

One major undertaking of this research was to determine the extent to which classification algorithms would use artifact activity to predict classes. To address this question, we separated EEG activity into ‘brain-only’ and ‘artifact-only’ using an objective ICA process: EEGlab’s *IClabel* algorithm. Furthermore, we had two baseline conditions to compare to: the EMG and ‘All ICs’. In an effort to make the process more robust to preprocessing bias, we also chose two separate approaches for our feature generation. Our returned classifier accuracy with both feature generation methods show a significant advantage for the ‘artifact-only’ condition versus the ‘brain-only’ condition. Furthermore, we found additional significant differences when using ‘All ICs’ and then again when using the EMG feature data. These group results can be seen in Figure 7, and participant level results can be found in the Appendix, Figure 3A.

**Figure 7.**
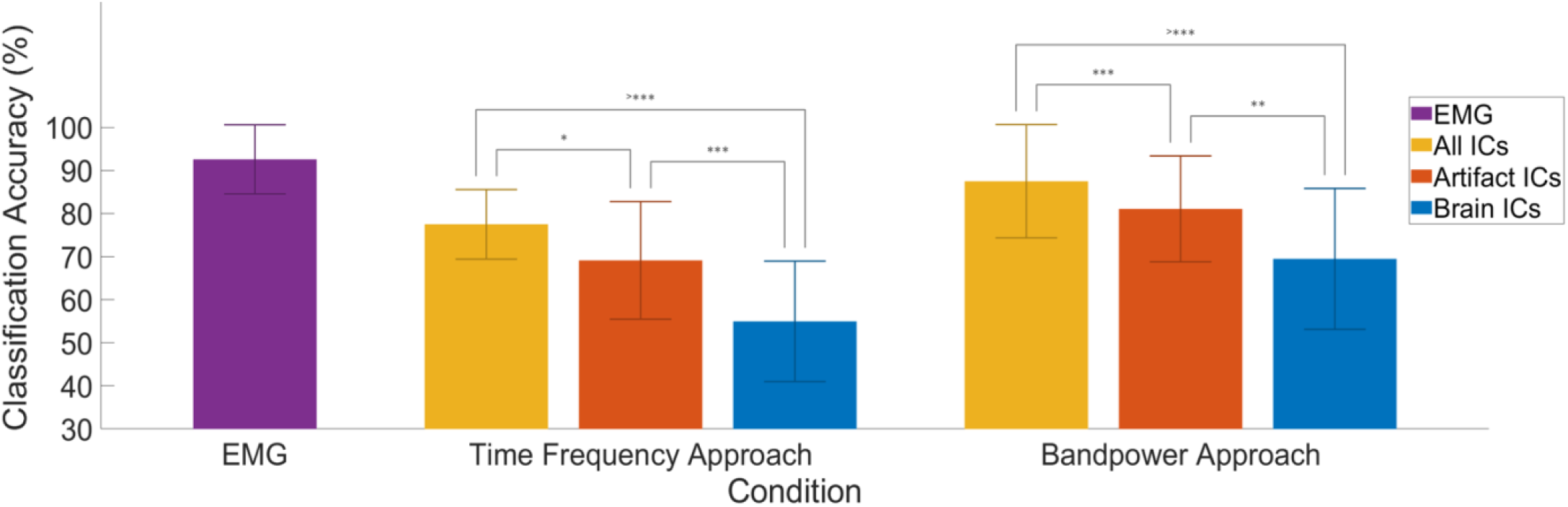
Comparison of Classifier Conditions. Group averaged classification accuracy is displayed for each of the separate conditions in a 5-class all-vs-one classification. “Brain” and “Artifact” consisted of 10 ICs across 10 bins of time, in 5 frequency bands, with dimensionality reduction to 70 features. “All ICs” was calculated by the same procedure, only with all ICs available. “EMG” used 7 channels on the side of movement, binned into 10 time bins each for features. Significance levels: * < 0.05, ** < 0.01, *** < 0.001

In addition to the overall condition comparison, we performed a subanalysis focusing on the differing frequency bands. Here, we observe that overall accuracy values are lower for the brain-only condition in both approaches, and that the theta band returns the highest accuracy in the artifact-only condition, while the gamma band returns the highest accuracy in the brain-only condition, as seen in Figure 8.

**Figure 8.**
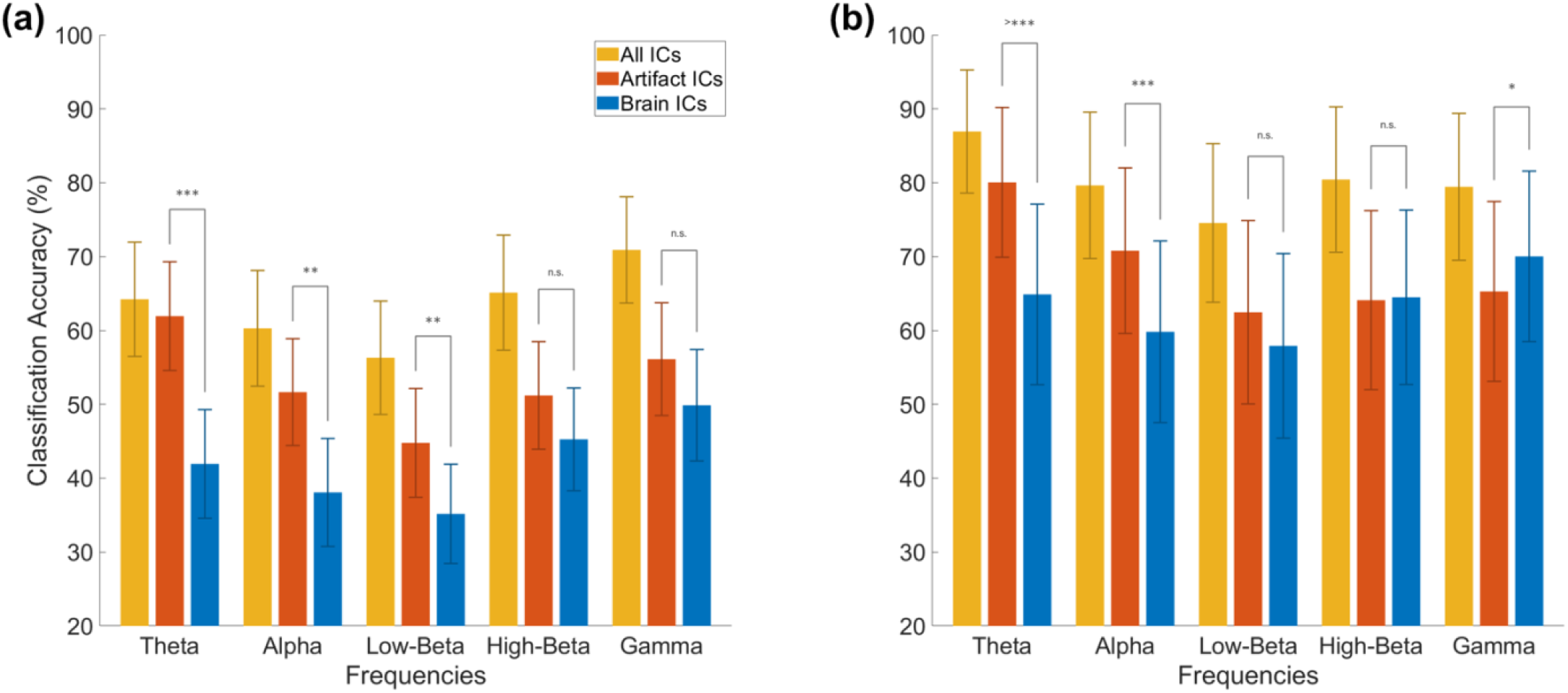
Frequency based analysis of EEG conditions. Results from the time frequency analysis approach (**a**) and bandpower (**b**) in each of 5 frequency bands. The artifact and brain conditions consisted of 10 ICs. The time frequency approach used 14 features for each frequency, the bandpower approach used 64 features. Significance levels: * < 0.05, ** < 0.01, *** < 0.001

### 3.5 Interpretation of Classifier Results

#### 3.5.1 Motivation

Beyond performing a simple comparison of artifact versus brain-derived activity with respect to classification accuracy, we also sought to understand which features were utilized by the classification algorithm in order to maximize class separability between classes. To do this, we interpreted the weight vectors produced within the TFA approach and re-projected them into sensor space. Additionally, in the bandpower approach, we averaged over each channel’s activity for all trials (in a given frequency band) to create topographies for each of the three conditions (all data, artifact, brain).

#### 3.5.2 Visualization of TFA approach

We interpreted the binary weight vectors returned from the all-vs-one classification algorithm by re-representing them in the sensor space using methods found in [26]. To begin, we first multiplied the weights with the coefficients saved in the PCA step of our pipeline, projecting back to a 500 dimension space (10 IC x 5 frequency band x 10 time bins), as seen for a single participant and single task in Figure 9.

**Figure 9.**
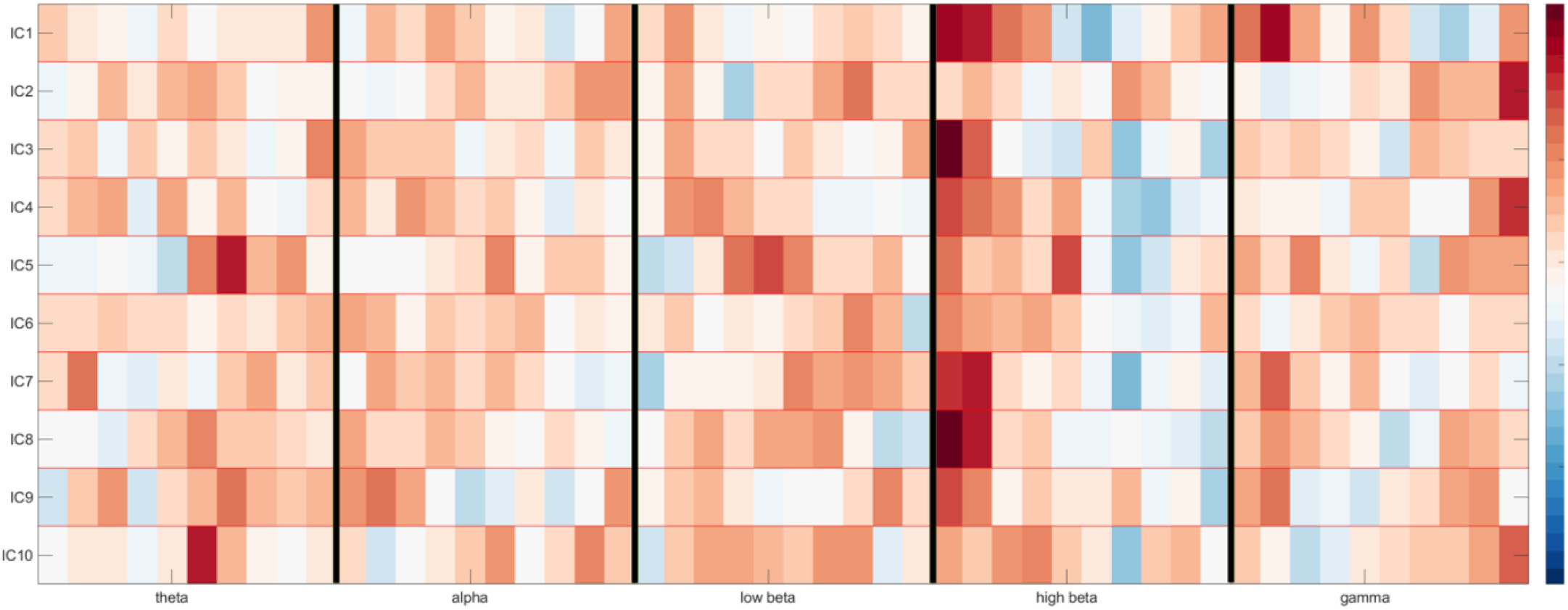
Reprojection of the time-frequency domain of 10 brain components as weighted by the classification model. The y-axis contains the ICs, whereas the x-axis is split into 10 time bins for each of the 5 chosen frequency bands, separated by vertical black lines. Maximum activations can be seen as red, minimum activations as blue.

This reprojection shows us the time-frequency domain of each of the 10 brain components as weighted by the classification model. If we then take the values from one of these frequency bands, and multiply them by the ICA matrix, we can visualize the topography, as seen for the alpha band in Figure 10 (top) below.

**Figure 10.**
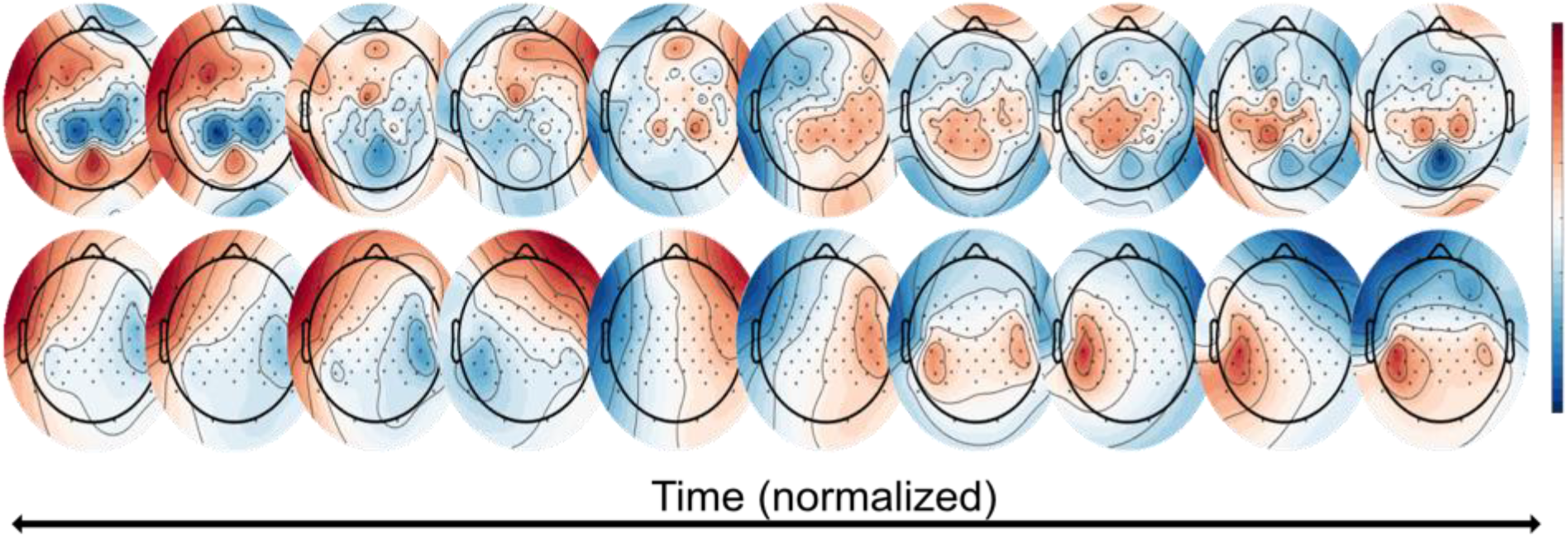
Classifier visualization of the TFA approach. (**Top**) These topographies represent the total ICA matrix multiplied with the first time bin of the TFA for the high beta frequency band. (**Bottom**) These topographies represent the further multiplication by the single task’s (here: “painting”) averaged covariance matrix. These topographies can be thought of as the “ideal” topography over time that maximizes the ability to separate the task from others with respect to the classification weight vectors. This patient used the left hand.

With this in mind, we then multiplied this by the covariance matrix of the epoched task data, allowing us to visualize the sensor space for that single task. These task-based topographies can be thought of as an “ideal” topography over time that maximizes the ability to separate the task (i.e. the “painting” task, where the participant moves the arm back-and-forth, here, the left arm) from the others with respect to classification accuracy weight vectors, as seen in Figure 10 (bottom).

#### 3.5.3 Visualization of Bandpower approach

The bandpower approach provided us with the opportunity to average across all trials for a given channel in a given frequency band, and then plot these results as a topography for each task. Doing so revealed to us activity localized in the frontal channels in the artifact condition, and over the motor cortex in the brain condition. However, the topographies also showed us that in the condition containing all available data, activity (in this case the high beta frequency) very closely mimics that of the artifact condition, as seen in Figure 11.

**Figure 11.**
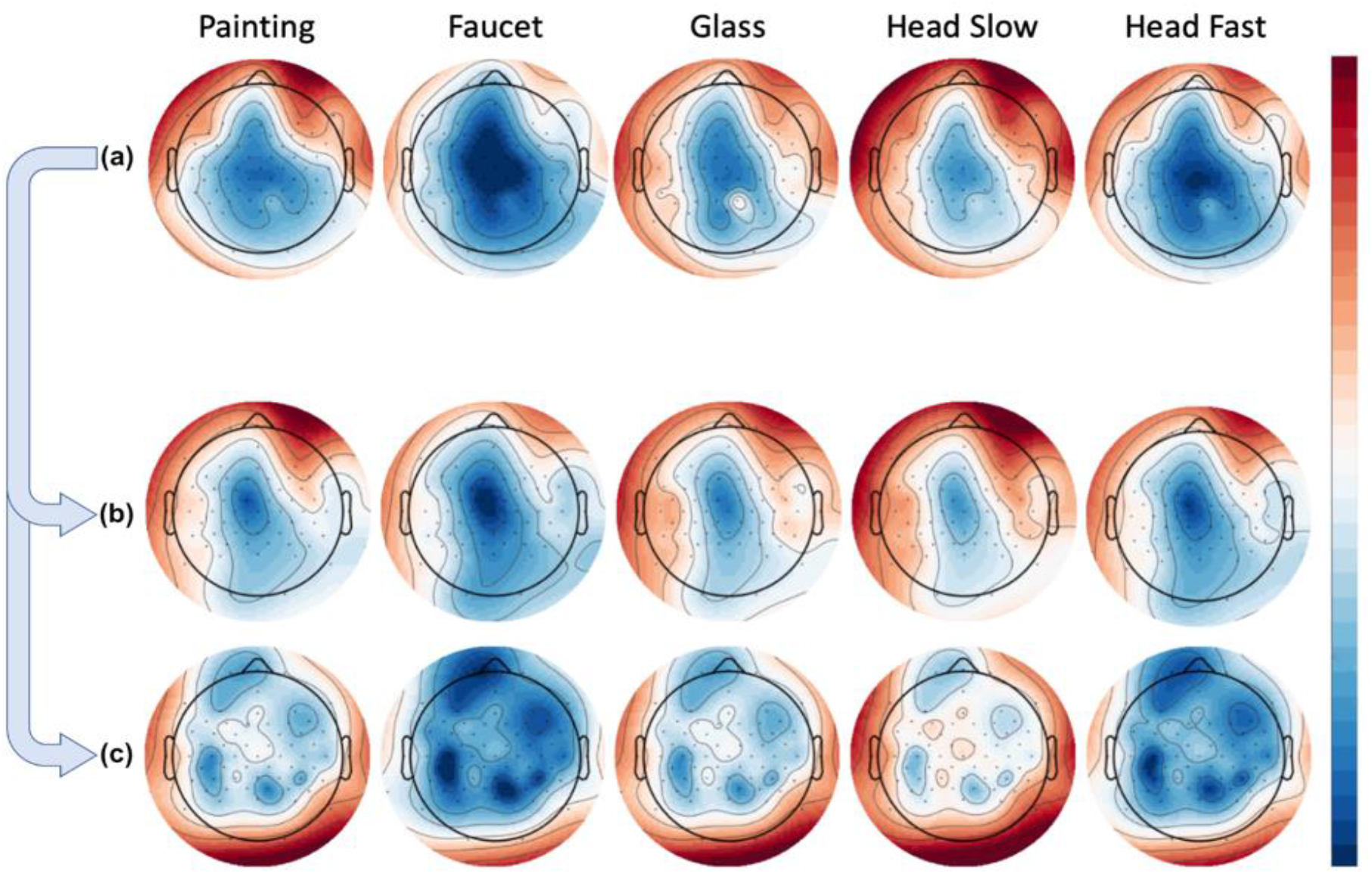
Group averaged bandpower per condition over high beta frequency. Rows represent tasks (Painting, Faucet, Glass, Head Slow, Head Fast). (**a**) Using all available data. (**b**) Artifact-only condition. (**c**) Brain-only condition.

### 3.6 Common Spatial Patterns (CSP) for Feature Selection

In an effort to further scrutinize the activity remaining within the brain-labeled ICs, we used CSP, a common state-of-the-art methodology that finds spatial patterns which maximize the variance between classes. Our rationale was that since artifacts have in general led to the greatest classification accuracy, if they remained somehow present in the data, this algorithm might be susceptible to utilizing the artifact-contaminated data in its components. In Figure 12, the subfigures on the left (x1) represent an instance of “cleaned” brain-activity from a patient, while on the right (x2), we see remaining artifacts despite the same pipeline from another patient. In detail, subfigures a1 and a2 show continuous EEG signal before (blue) and after (red) applying ICA for artifact removal. The top 10 brain-labeled ICs are found in subfigures b1 and b2, and top 4 spatial projections of the CSP algorithm are visualized in c1 and c2. It appears that artifact-contaminated channels could be successfully cleaned with the ICA approach, as the top 10 brain-ICs from both patients show topographies typical of brain activity. Critically, however, after the CSP step, we see from the projections that artifact removal was not entirely successful for the patient’s data as visualized in c2. In particular, the first CSP component, which is the projection with the highest variance, has a topography typically seen from eye movement artifact.

**Figure 12.**
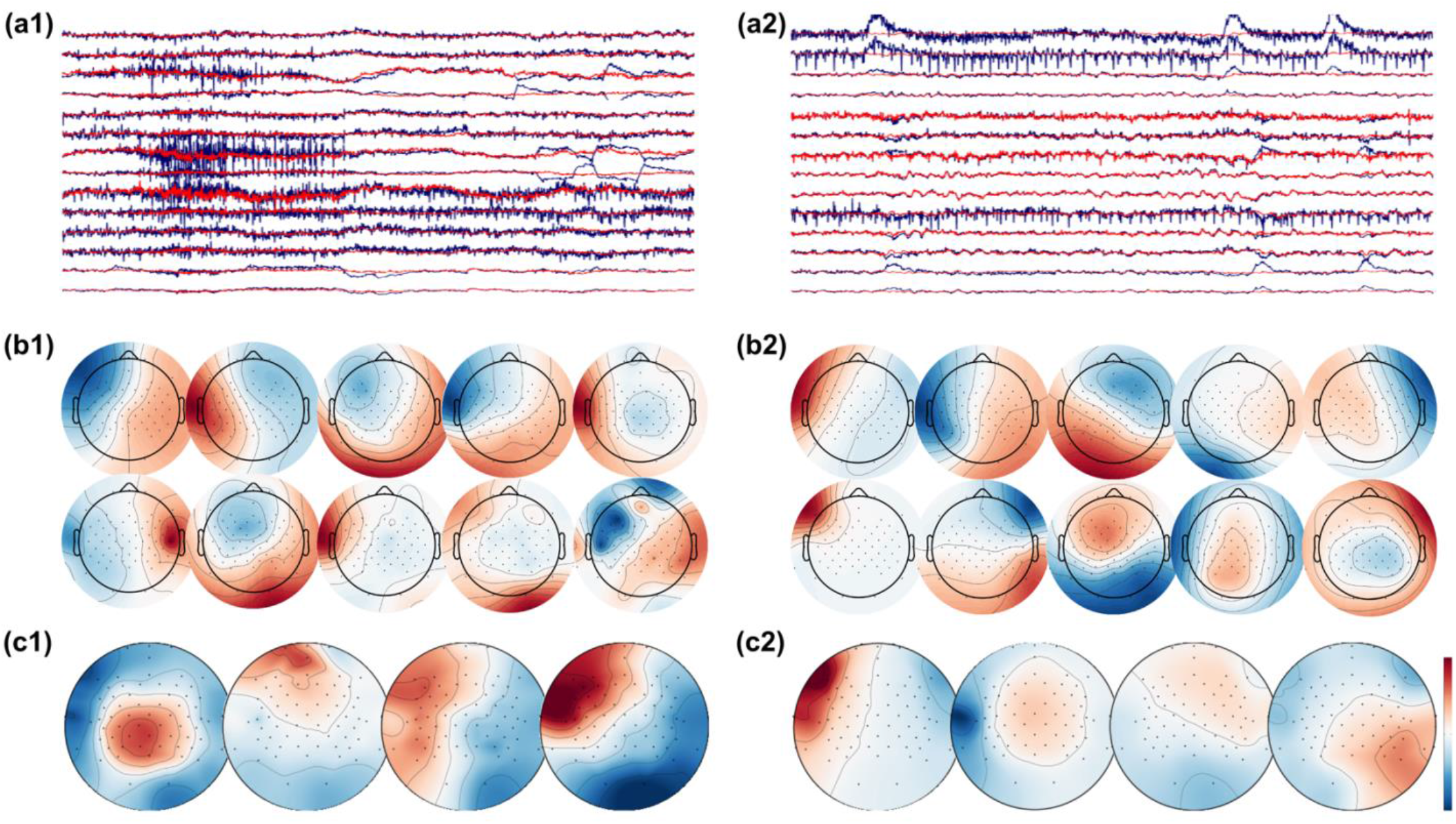
EEG signals before (raw EEG, blue) and after (clean EEG, red) artifact removal with ICA from patient 1 (**a1**) and patient 2 (**a2**). The top-10 brain ICs are shown in (**b1**) and (**b2**), and the top-4 spatial pattern projections after applying the CSP algorithm shown in (**c1**) and (**c2**). The topographies show successful removal of artifacts in patient 1, while artifacts are present in the topographies of patient 2.

## 4. Discussion

### 4.1 Implications

As EEG-based BCIs are becoming increasingly relevant in research, clinical, and consumer applications, it also becomes increasingly important to understand the origin of the signal features which underlie the utilized output, especially in scenarios where the BCI is designed to identify a physiological brain state. As the results of this study show, state-of-the-art automated EEG cleaning pipelines using ICA are an effective method to remove EEG artifacts from data recorded from patients with stroke wearing a VR headset and performing naturalistic movements in a neurorehabilitation setting. Across two different feature extraction methods, classifier visualization shows that topographies consistent with brain-activity are recovered from the cleaned data as the most informative features.

On the other hand, when the EEG data is not cleaned, the most informative features show artifact topographies in both feature extraction methods. This pattern can also be seen in the classification accuracy when comparing the brain-signal vs. the artifact portion of the EEG data, as artifact components are consistently more informative and are therefore selected by a classifier if available. Furthermore, even after the cleaning process, it is possible to “recover” remaining artifact information in some subjects when using methods that further reduce dimensionality based on class separability, such as CSP. An example of this can be seen in Figure 12, where from the previously cleaned data, one CSP extracts brain-signals and one CSP extracts artifacts.

This finding has significant therapeutic relevance for BCI-based neurorehabilitation where the goal is to use brain activity to induce neural plasticity at the circuit level [17]. The results of this study highlights the risk of inadvertently using artifact signal components to inform the therapy, which would likely not only lead to a reduced therapeutic effect, but would also be difficult to detect, as the BCI appears to function adequately. Moreover, sophisticated machine learning approaches to improve the BCI’s classification performance such as CSP can actually be counterproductive from the point of view of a BCI-based motor neurorehabilitation paradigm.

### 4.2 Limitations

We consider a limitation of this study to be the generalizability of this main result beyond the scenario of decoding EEG-signals during naturalistic movements. In our study, the different neurophysiological states of interest are correlated with different motor trajectories. This of course makes artifacts especially informative and therefore problematic, but this is precisely the problem in BCI-based motor neurorehabilitation paradigms. Though our study also relies on the separability of EEG data into an “artifact” part and a “brain signal” part using ICA, a complete separation is not possible with real data. Additionally, whereas we have balanced the number of ICA components, it is still possible that this overlap is asymmetric, with more cross contamination coming from one class than the other. Nevertheless, ICA is a standard approach for EEG cleaning and we consider this the most relevant method in practice.

### 4.3 Conclusion

This study highlights the need to consider the influence of movement-related artifacts when designing BCI-based neurorehabilitation paradigms to detect neurophysiological brain states. Standard EEG cleaning methods with ICA can be used to aggressively remove noise-related components, if a sufficient number of EEG channels are available. When using machine-learning approaches to analyze the data, we suggest visualizing the spectra and topographies of the most informative features used by the classifiers as a matter of best practice. Finally, when decoding physiological brain states for therapeutic applications, feature extraction should be informed by physiology, rather than automatically optimized to maximize classification accuracy.

## Author Contributions

Conceptualization, E.J.M, U.Z., and C.Z.; methodology, E.J.M., P.R., and C.Z.; software, E.J.M., P.R., S.K., and C.Z.; validation, E.J.M. and P.R.; formal analysis, E.J.M., P.R., and C.Z.; investigation, E.J.M., P.R., and C.Z.; resources, E.J.M., P.R., S.K., U.Z., and C.Z.; data curation, E.J.M. and P.R.; writing—original draft preparation, E.J.M.; writing—review and editing, E.J.M, P.R., and C.Z.; visualization, E.J.M.; supervision, U.Z. and C.Z.; project administration, E.J.M, U.Z., and C.Z.; funding acquisition, C.Z. and U.Z. All authors have read and agreed to the published version of the manuscript.

## Institutional Review Board Statement

The study was conducted according to the guidelines of the Declaration of Helsinki, and approved by the Ethics Review Committee of the Medical Faculty of Eberhard Karls University Tübingen (Protocol BNP-2019-11).

## Informed Consent Statement

Informed consent was obtained from all subjects involved in the study.

## Data Availability Statement

Data is available upon request.

## Acknowledgments

We thank Mareike Spieker for her assistance in data collection.

## Appendix

**Figure 1A:**
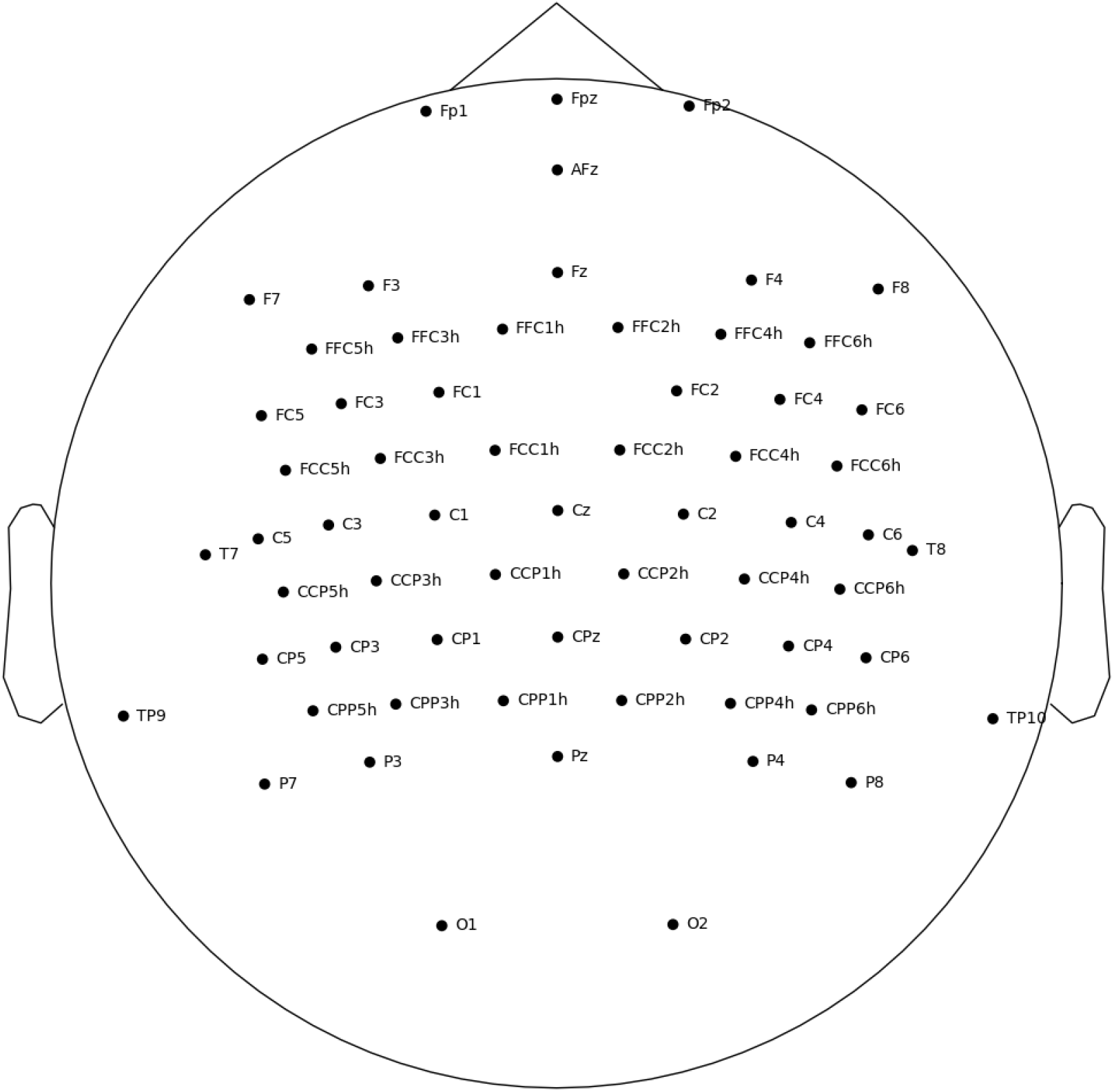
EasyCap 64 channel layout, with additional electrodes are over the motor cortex.

**Figure 2A:**
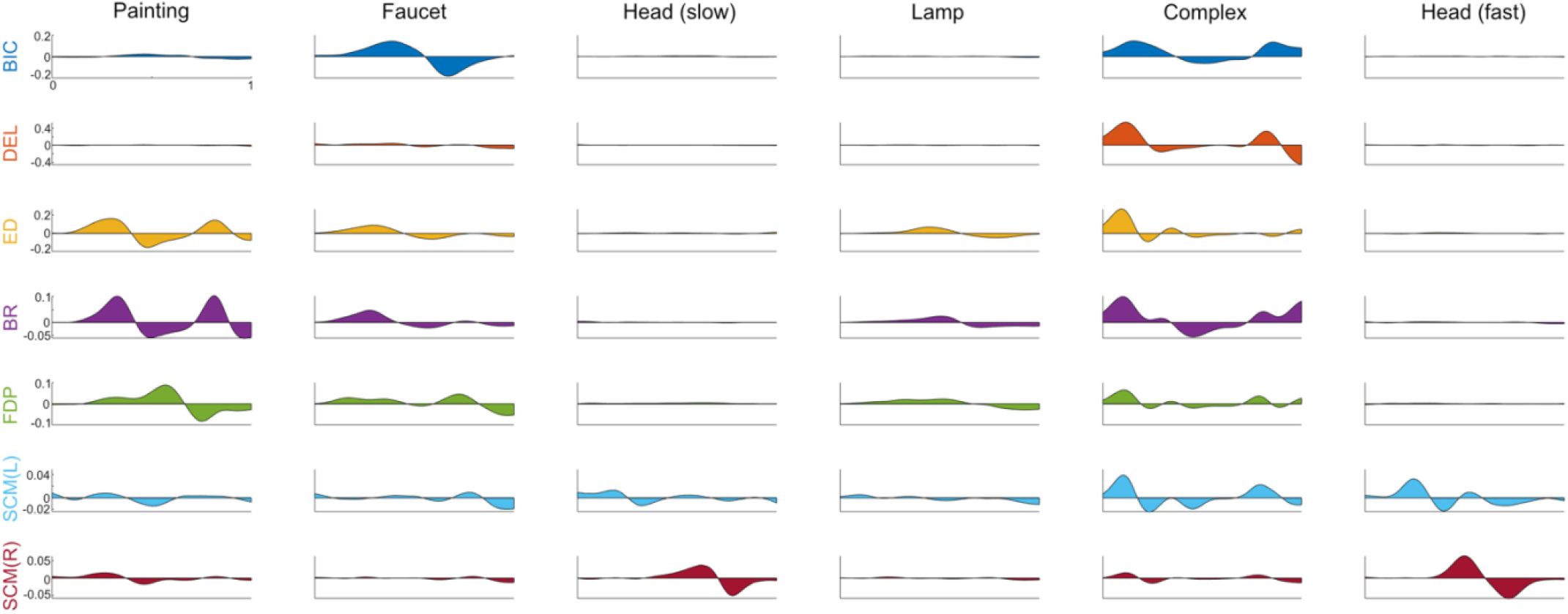
Complete EMG traces. Rate of change of muscle activation during the task recorded from 7 different EMG channels, by task, averaged across participants, task duration normalized. Each column represents a task: painting, faucet, head (slow), lamp, complex, head (fast). Each row represents a bipolar EMG channel, recorded from the following muscles: Biceps, Deltoid, Extensor Digitorum, Flexor Digitorum Profundus, Sternocleidomastoid (left, right).

**Figure 3A:**
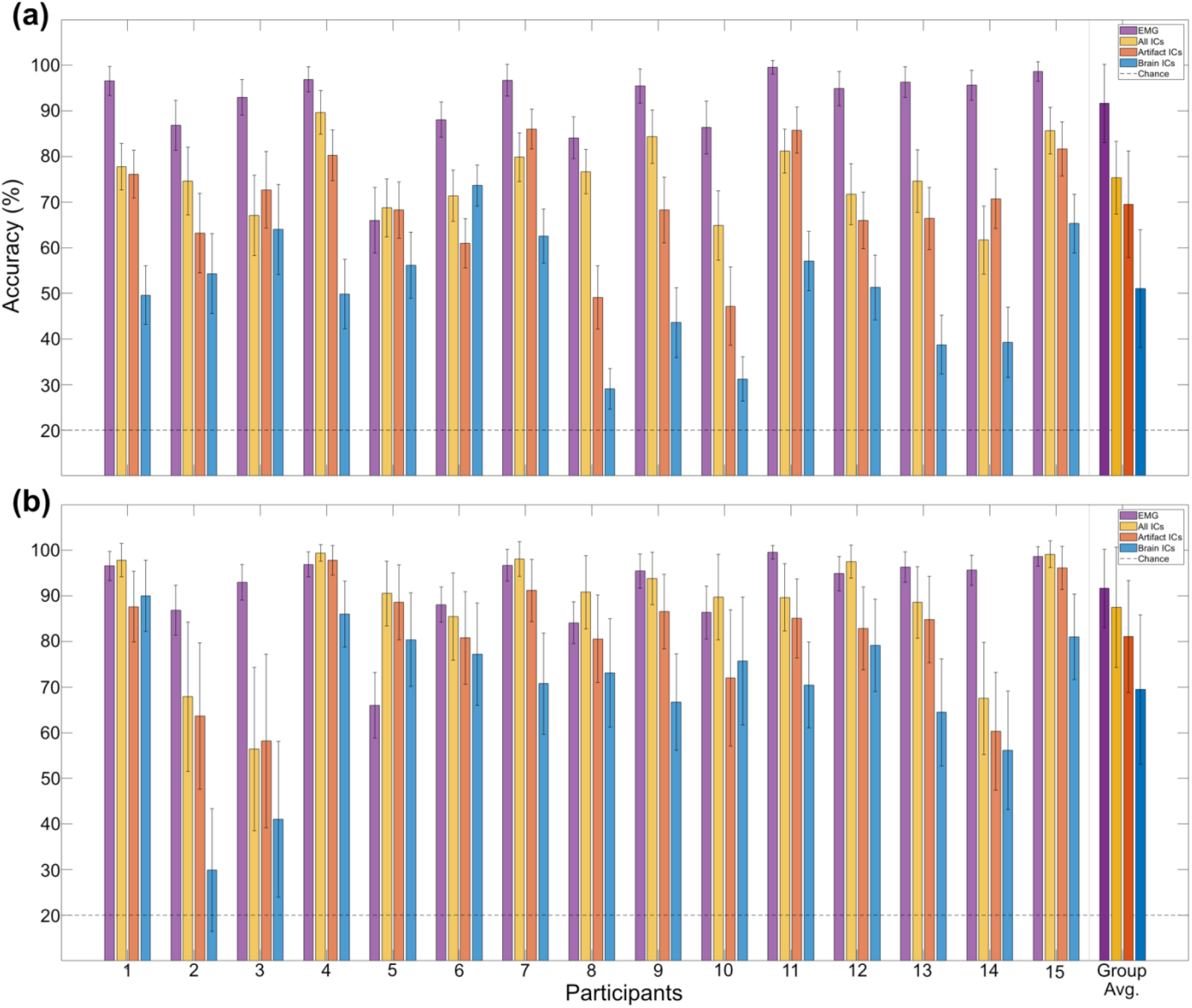
Participant-level Classification Accuracy. Displayed from Time-Frequency Analysis (**a**) and Bandpower Analysis (**b**), also compared with EMG. Multi-class classification between 5 equally likely movements, chance level indicated with dotted line

## Footnotes

1 https://vive.com/

2 https://store.steampowered.com/valveindex/

3 https://www.unrealengine.com/

4 https://www.mathworks.com/products/matlab.html

5 https://mne.tools/stable/index.html

6 https://www.mathworks.com/help/stats/fitcecoc.html

7 https://www.python.org/

8 https://scikit-learn.org/stable/

